# Direct Visualization of Translesion DNA Synthesis Polymerase IV at the Replisome

**DOI:** 10.1101/2022.04.23.489275

**Authors:** Pham Minh Tuan, Neville Gilhooly, Kenneth J. Marians, Stephen C. Kowalczykowski

## Abstract

In bacterial cells, DNA damage tolerance is manifested by the action of translesion DNA polymerases that can synthesize DNA across template lesions that typically block the replicative DNA polymerase III. It has been suggested that one of these TLS DNA polymerases, DNA polymerase IV, can either act in concert with the replisome, switching places on the β sliding clamp with DNA polymerase III to bypass the template damage, or act subsequent to the replisome skipping over the template lesion in the gap in nascent DNA left behind as the replisome continues downstream. Evidence exists in support of both mechanisms. Using single-molecule analyses we show that DNA polymerase IV associates with the replisome in a concentration-dependent manner and remains associated over long stretches of replication fork progression under unstressed conditions. This association slows the replisome, requires DNA polymerase IV binding to the β clamp but not its catalytic activity, and is reinforced by the presence of the γ subunit of the β clamp-loading DnaX complex in the DNA polymerase III holoenzyme. Thus, DNA damage is not required for association of DNA polymerase IV with the replisome. We suggest that under stress conditions such as induction of the SOS response, the association of DNA polymerase IV with the replisome provides both a surveillance/bypass mechanism and a means to slow replication fork progression, thereby reducing the frequency of collisions with template damage and the overall mutagenic potential.

**Significance:** Damage to the nucleotide bases that make up the DNA in chromosomes creates a problem for their subsequent accurate duplication each time a cell divides. Typically, the cellular enzymatic machinery that replicates the DNA cannot copy a damaged base and specialized trans-lesion DNA polymerases, which are prone to making errors that result in mutations, are required to copy the damaged base, allowing replication to proceed. We demonstrate that the bacterial replisome, which is comprised of the enzymes required to replicate the chromosome, can associate with one of these specialized trans-lesion polymerases over long distances of replicated DNA. This association slows the speed of replication, thereby reducing the chance of mutations arising in the cell under conditions of stress.

## Introduction

Replication fork progression in E. coli is catalyzed by a replisome comprised of the DNA polymerase III holoenzyme (Pol III HE), which provides both the leading- and lagging-strand DNA polymerases, the hexameric replicative DNA helicase, DnaB, and the Okazaki fragment primase, DnaG (1). The Pol III HE itself contains ten subunits: two copies of the core DNA polymerase (αεθ, where α is the catalytic DNA polymerase and ε is the proofreading 3’→5’ exonuclease), the dimeric sliding processivity clamp β, and the DnaX complex, τ_2_γδδ’χψ, which loads β to the primer template (1). The core polymerases are bound to the DnaX complex via interaction with the τ subunit (2, 3). An alternate form of the DnaX complex, τ_3_δδ’χψ, allows the assembly in vitro of a Pol III HE with three core polymerases (4), the existence of which in vivo is supported by imaging of fluorescently tagged polymerase subunits (5, 6). However, cells that do not produce the γ subunit are UV-sensitive and have reduced mutagenic break repair (7), an activity that requires DNA polymerase IV (Pol IV) (8).

DNA synthesis catalyzed by the Pol III HE is highly accurate, with an error rate of roughly 10^-7^ (9). In general, Pol III cannot bypass bulky template lesions, although we have shown that it can bypass a cis-syn thymidine dimer (10). Bypass of most template lesions in the cell is ascribed to the action of translesion DNA synthesis (TLS) polymerases of which E. coli has three: DNA polymerases II, IV, and V (11). These polymerases demonstrate different activities with various template lesions, with Pol V Mut, a RecA-activated form of Pol V (12, 13) [UmuD’_2_UmuC (14, 15)], being the major activity under conditions of high replication stress when the SOS response has been activated (16, 17). Pol IV, which is encoded by *dinB* (18), is a Y family DNA polymerase that is well conserved from bacteria to eukaryotic cells. Pol IV has long been thought to be present at the highest concentration of all DNA polymerases in E. coli under unstressed conditions, about 250 copies per cell, with this concentration increasing about 10-fold upon induction of the SOS response (19). Interestingly, a recent study that measured the signal generated by fluorescently-tagged Pol IV molecules in live cells argued that these high values were inaccurate, with the basal level of Pol IV being about twenty copies/cell and the SOS-induced level about 280 copies/cell (20).

Unlike Pol III HE, which is a rapid and highly processive DNA polymerase (21–23), Pol IV is distributive, incorporating only one nucleotide per primer binding event (24). However, like all E. coli DNA polymerases, it can interact with the β processivity clamp and in so doing its processivity increases to 300-400 nucleotides (24). Overproduction of Pol IV in the absence of replication stress slows DNA replication (25, 26) and it has been demonstrated that it is the main factor contributing to slowing replication fork progression under conditions of stress (27). Studies in vitro have shown that Pol II and Pol IV can generate slow-moving replication forks in the presence of DnaB and DnaG and that very high concentrations of Pol IV can slow the canonical Pol III-DnaB-DnaG replisome (28).

The ability of all E. coli TLS polymerases to bind to β led to the formulation of the “tool belt” model to account for rapid localization of TLS polymerases to the site of replisome stalling (29) with the concept being that because β is a dimer, the TLS polymerase could ride along with the replisome on the same sliding clamp that was bound to the α subunit of the Pol III HE and switch places with a stalled Pol III to catalyze lesion bypass. How the polymerase switch occurs is still unclear. It has been suggested that switching only occurs at a stalled polymerase and that the stalled Pol III dissociates from β and is replaced by Pol IV (28, 30, 31). It has also been a common view that Pol IV association with the replisome is concentration-dependent (32, 33), accounting for association only when it is needed, i.e., when the SOS response is induced. We have shown that Pol IV-dependent bypass at a thymidine dimer in the leading-strand template in the presence of an active replisome competes with lesion skipping (34), when the stalled leading-strand polymerase cycles ahead to a new primer made on the leading-strand template to continue replication downstream (35), suggesting that polymerase switching had occurred.

Here we have addressed association of Pol IV with the replisome using single-molecule DNA replication (23). We show that in the absence of template damage Pol IV can associate in a concentration-dependent manner with the replisome and proceeds along with it during replication fork progression. Association of Pol IV with the replisome requires its β binding motif, is stabilized by the presence of the γ subunit of the DnaX complex, and generates two classes of replisomes: those without Pol IV bound that progress rapidly and those with Pol IV bound that proceed slowly. Slowing of the replisome by Pol IV does not require its catalytic activity, suggesting that the decrease in rapid replication fork progression is a result of Pol IV binding directly to one of the two polymerase binding clefts on β. The constant presence of Pol IV in the replisome may act as a template damage surveillance mechanism.

## Results

### Pol IV slows progression of the replication fork

Rolling circle DNA replication in a flow cell was visualized by single-molecule TIRF microscopy (23). Tailed form II DNA templates were anchored to a biotinylated glass surface via the biotin-streptavidin interaction and any free DNA templates were washed out of the flow cell (Figures 1A and S1). DNA replication was performed in two steps: 1) Replisome assembly reactions contained DnaB, DnaC810 (which loads DnaB to the template (36)), Pol III HE, three dNTPs, four NTPs, and Pol IV as indicated (“Assembly”, Figure 1A). In this step the replisome forms on the template and idles because of the missing dNTP. It is important to note that all proteins not associated with the replisome on the template are then washed out of the flow cell. This is the only stage of the reaction where Pol IV is present; thus, the reaction is single-turnover with regard to Pol IV. Replication was initiated in the “Start Reaction” by the introduction of SSB, β, DnaG, all four dNTPs, and NTPs into the flow cell. Synthesized DNA is stained with SYTOX Orange, and the DNA visualized continuously by TIRF microscopy (Figures 1B and S1). A representative frame from a typical movie (Movie S1) of a reaction in the absence of Pol IV and a kymograph of a typical molecule are shown in Figures 1B and 1C, respectively. Trajectories of replicating molecules are fitted to a three-segment line (Figure 1D) to determine the start and end points of DNA replication. The rate of replication fork progression is determined by dividing the length of DNA synthesized by the time elapsed from initiation to termination of DNA synthesis. Processivity of DNA replication is given by the length of DNA synthesized.

**Figure 1:**
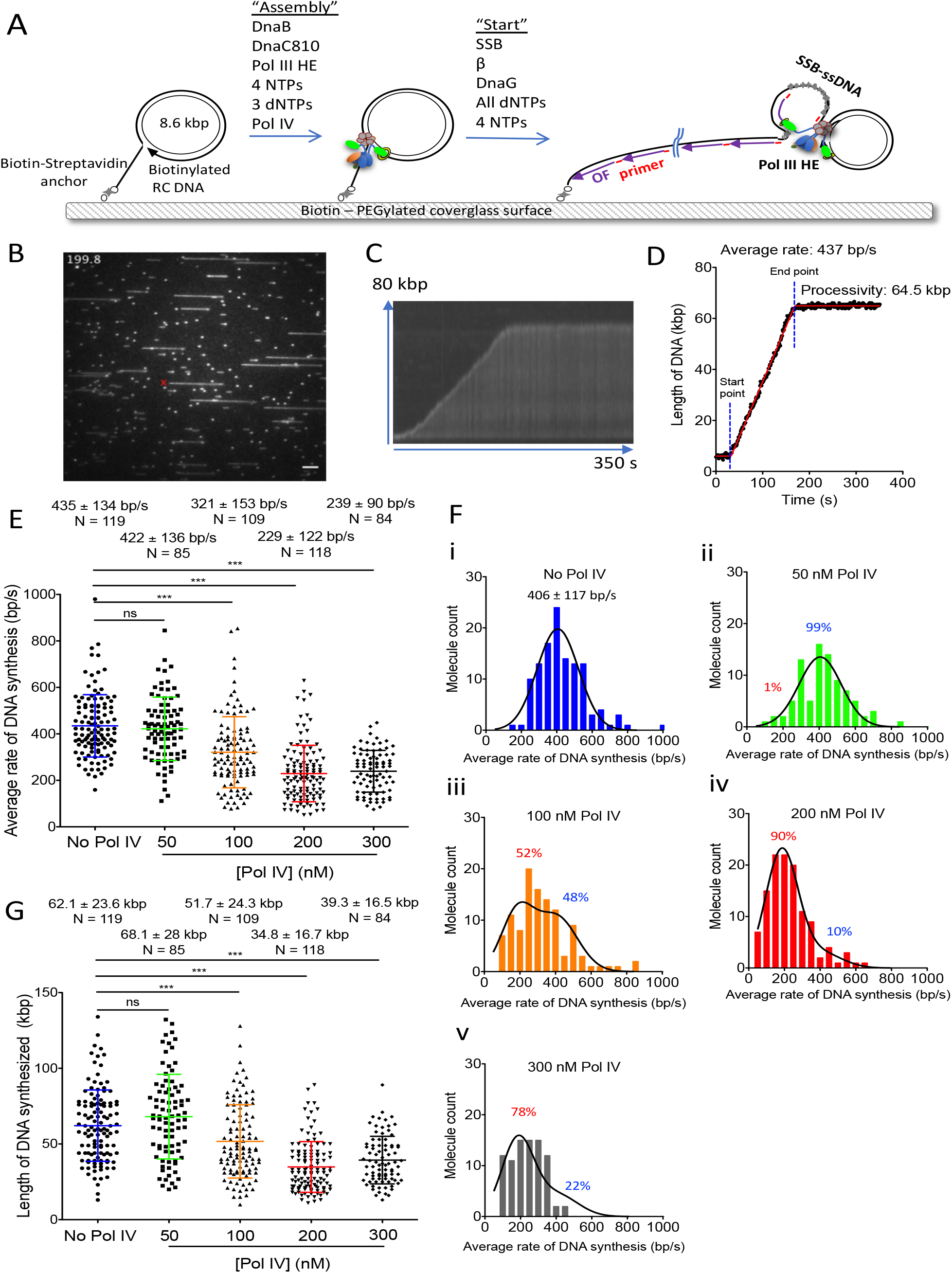
Pol IV slows the rate of replication fork progression of Pol III HE replisomes in a concentration-dependent manner. (A) Schematic of the rolling circle DNA replication assay. (B) Micrograph showing a typical frame from a reaction in the absence of Pol IV. Scale bar, 5 μm, equivalent to 17.5 kb double-stranded DNA at a flow rate of 1250 μl/h. (C) Kymograph and (D) tracking trajectory of a representative molecule with a three-segment line fit (red). The start and end points of replication fork progression are indicated by the dotted blue dot lines. (E) Average rates of DNA synthesis from the live imaging reactions in the presence or absence of the indicated concentrations of Pol IV in the assembly step. (F) Distribution of the average rates of DNA synthesis for the data shown in panel E. Panels (i), (ii), (iii), (iv), and (v) are for no Pol IV, 50 nM Pol IV, 100 nM Pol IV, 200 nM Pol IV, and 300 nM Pol IV, respectively. The black curve in (i) is a single Gaussian fitting with the indicated mean value and standard deviation (SD). The black curves in (ii)-(v) show the sum of two Gaussian curves using values constrained to 406 bp/s ±117 bp/s and 185 bp/s ±90 bp/s for the fast and slow components, respectively. (G) Processivities of replication forks from the live imaging reactions in the presence or absence of the indicated concentrations of Pol IV in the assembly step. Mean values are given with standard deviations. (*) and (***) denote p < 0.05 and p < 0.0001, respectively, determined by the student’s t-test. The results represent the average of more than ten independent experiments at each condition. N, molecules.

The addition of increasing concentrations of Pol IV in the “Assembly” step progressively reduced the average speed of replication forks from 435 bp/s in the absence of Pol IV to roughly half that in the presence of 200-300 nM Pol IV (Figure 1E; Movie S2 shows an example of replicating molecules formed in the presence of 200 nM Pol IV). In the absence of Pol IV, the replication fork rates can be adequately fit by a broad single-Gaussian distribution (406 ±117 bp/s; Figure 1F-i), as was reported previously (23). However, in the presence of increasing amounts of Pol IV, the replication fork rate distributions showed both broadening and a shift of the population to slower rates. This behavior suggested the presence of potentially two Gaussian distributions, and analyses showed, particularly at 100-200 nM Pol IV, that the distribution was best described as the sum of two Gaussian distributions: one with the speed of replication forks formed in the absence of Pol IV, and one with a slower average rate of about 185 +/-90 bp/s (Figure 1F). The change in the relative compositions (amplitudes) of the two Gaussian components as a function of Pol IV concentration suggests an apparent Kd of ~80-100 nM for Pol IV binding to the replisome. At 200-300 nM Pol IV most of the replicating molecules were of the slow class. The speed of slowly moving molecules did not change during the measured trajectories, suggesting that Pol IV remained bound to the replisome for the duration of DNA synthesis; had the Pol IV dissociated at any time, we expected that the rates of the replisomes would have increased to the values observed in the absence of Pol IV. The average processivity of replication forks decreased similarly as a function of Pol IV concentration from 62 kbp to about 35 kbp (Figure 1G). The efficiency of initiation of DNA replication (measured as the fraction of templates replicating in the overall field of view) also decreased as a function of Pol IV concentration (Table 1). These data suggest that when Pol IV was present at the time of assembly with the replisome, two populations were created: one where the replisome had an associated Pol IV molecule (slow) and another where no Pol IV was associated (fast).

**Table 1:** Effect of Pol IV on initiation, replication fork progression, and processivity of replisomes formed with either the DiPol III holoenzyme or TriPol III holoenzyme

| Pol III HE | Reaction | Average rate (bp/s) | Processivity (kb) | DNA template initiated (%) |
| --- | --- | --- | --- | --- |
| **DiPol** | No Pol IV | 435 ± 134 | 62.1 ± 23.6 | 15.9 |
|  | 50 nM Pol IV | 422 ± 136 | 68.1 ± 28.0 | 8.2 |
|  | 100 nM Pol IV | 321 ± 153 | 51.7 ± 24.3 | 8.6 |
|  | 200 nM Pol IV | 229 ± 122 | 34.8 ± 16.7 | 6.7 |
|  | 300 nM Pol IV | 248 ± 92 | 39.3 ± 14.6 | 4.1 |
|  | 400 nM Pol IV | 201 ± 90 | 25.2 ± 6.0 | 1.1 |
|  | 500 nM Pol IV | N.D. | N.D. | 0.2 |
|  | 200 nM Pol IV D8A | 251 ± 97 | 40.3 ± 18.4 | 5.4 |
|  | 500 nM Pol IV D8A | N.D. | N.D. | 0 |
|  | 200 nM Pol IV ΔC5 | 415 ± 131 | 62.7 ± 26.2 | 12 |
| **TriPol** | No Pol IV | 416 ± 194 | 62 ± 33.9 | 16.4 |
|  | 200 nM Pol IV | 339 ± 126 | 56.5 ± 27 | 13.9 |
|  | 500 nM Pol IV | 282 ± 161 | 40.4 ± 24.8 | 6.5 |

N.D. not detectable

### The presence of Pol IV in the replisome affects the burst rates of DNA synthesis

Our previous studies of DNA replication by single-molecule analyses demonstrated that replication fork progression occurred in a random series of bursts and pauses (23). DNA synthesis burst rates varied between 100 bp/s and 1000 bp/s, whereas pause times varied between 8 s and 25 s. We therefore asked whether Pol IV was affecting either DNA synthesis burst rates or pause times.

To analyze burst rates and pause times, replication fork trajectories were fitted with multisegment lines that divide the trajectory into segments of active DNA synthesis and inactive pauses (Figures 2A and 2B). Similar to the effect of Pol IV on the average rate of replication fork progression, increasing concentrations of Pol IV progressively decreased the average burst rate of DNA synthesis from 451 bp/s in the absence of Pol IV to about 280 bp/s in the presence of 200-300 nM Pol IV (Figure 2C). On the other hand, there was no effect of Pol IV on the average pause times (Figure 2D), neither duration nor number. Thus, the action of Pol IV to slow replication fork progression occurs during active polymerization.

**Figure 2:**
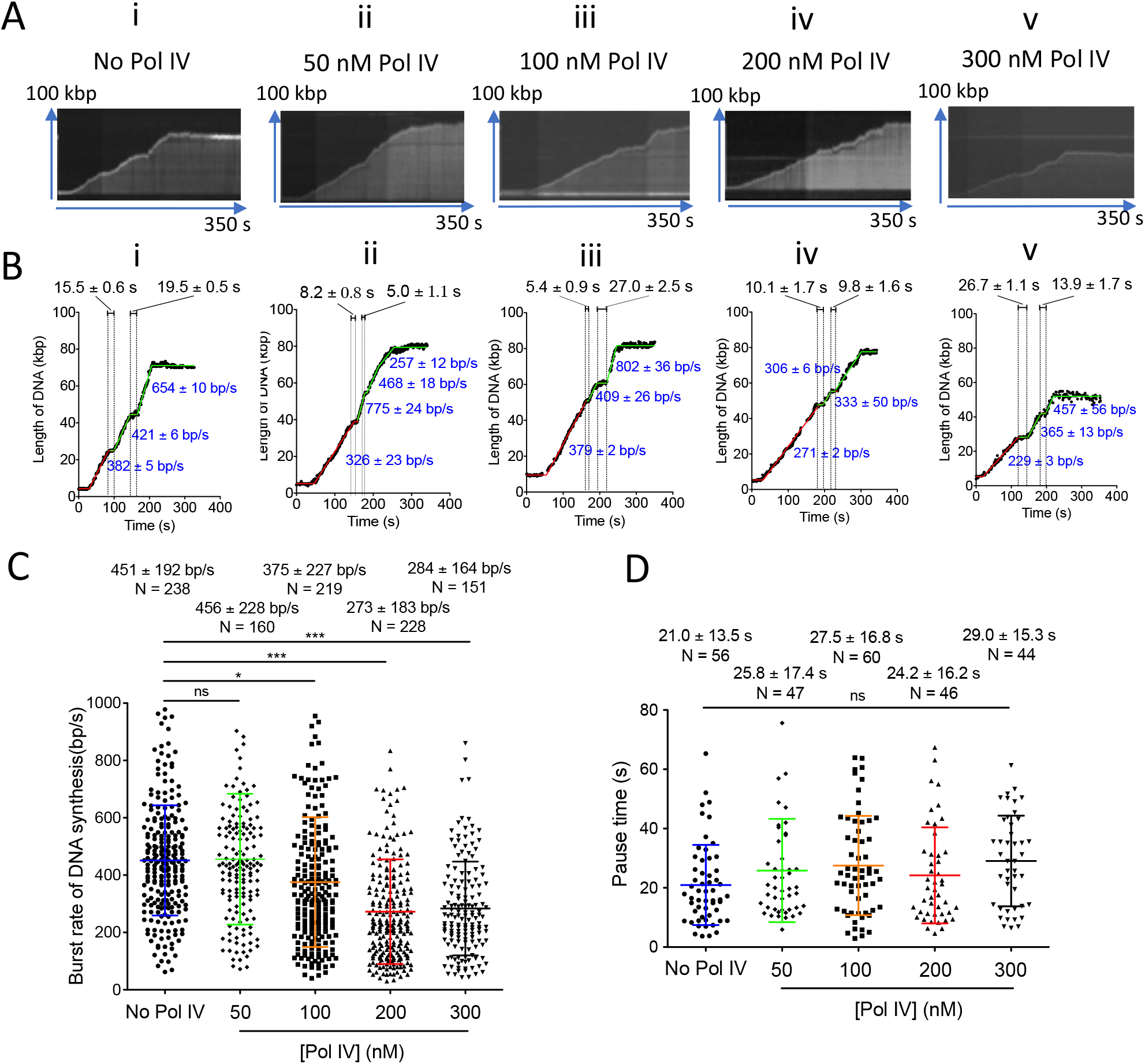
The presence of Pol IV in the replisome affects the burst rate of replication fork progression. (A) Typical kymographs and (B) tracking trajectories. (i), (ii), (iii), (iv), and (v), no Pol IV, 50 nM Pol IV, 100 nM Pol IV, 200 nM Pol IV, and 300 nM Pol IV, present in the assembly step, respectively. The trajectories were fit to multiple and overlapped fitting segments shown as green and red lines. The segment fittings give the burst rates of DNA synthesis and pause times as described under Experimental Procedures. (C) Average burst rates of DNA synthesis in the presence or absence of the indicated concentrations of Pol IV in the assembly step. (D) Average pause times during replication fork progression in the presence or absence of the indicated concentrations of Pol IV in the assembly step. Mean values are given with standard deviations. Mean values are given with standard deviations. (*) and (***) denote p < 0.05 and p < 0.0001, respectively, determined by the student’s t-test. N, molecules.

### Pol IV localizes with the active replisome

Given that Pol IV was only present in our replication reactions during the initial replisome assembly stage when there is no possibility of replisome advancement and was then subsequently washed out of the flow cell before the initiation of DNA replication, it seemed likely that it had to be associated with the moving replisomes. To address this question directly, we labeled Pol IV with the fluorescent dye Cy5 and examined its behavior in real time vis-à-vis the moving replisome. Labeled Pol IV retained 90% of the activity of the unlabeled wild type (Figure S2).

When 200 nM Cy5-Pol IV was present in the “Assembly” step and subsequently washed out before initiation of replication, the Cy5 signal migrated with the end of the replicating DNA where the template is located (Figure 3A and Movie S3). The kymographs, trajectories, and rates of movement of the DNA and the Cy5 signals were identical, with the distance between the two signals remaining constant throughout the course of replication fork progression (Figures 3A and S3D). Comigration was evident for most (~80%) of the replisomes (Movie S3). Replication fork rates fell into two classes: the forks with a bound Cy5-Pol IV presented with an average rate of 310+/-137 bp/s, whereas the forks lacking Cy5-Pol IV presented with an average rate of 440+/-115 bp/s (Figure 3B). The fraction of fork rates falling into the slow class is consistent with Pol IV occupancy at replisomes being 80-90% at this concentration, as in Fig 1F, iv. Furthermore, the Pol IV and DNA signals co-translocated for an average of at least 22 kbp (Figure 3C). This co-translocation distance is an underestimate because in some cases the fluorescent signal disappears, most likely because of photobleaching of the Cy5 fluorophore. Taken together, these data suggest that the labeled Pol IV binds to the replisome during assembly and rides along with it during replication fork progression.

**Figure 3:**
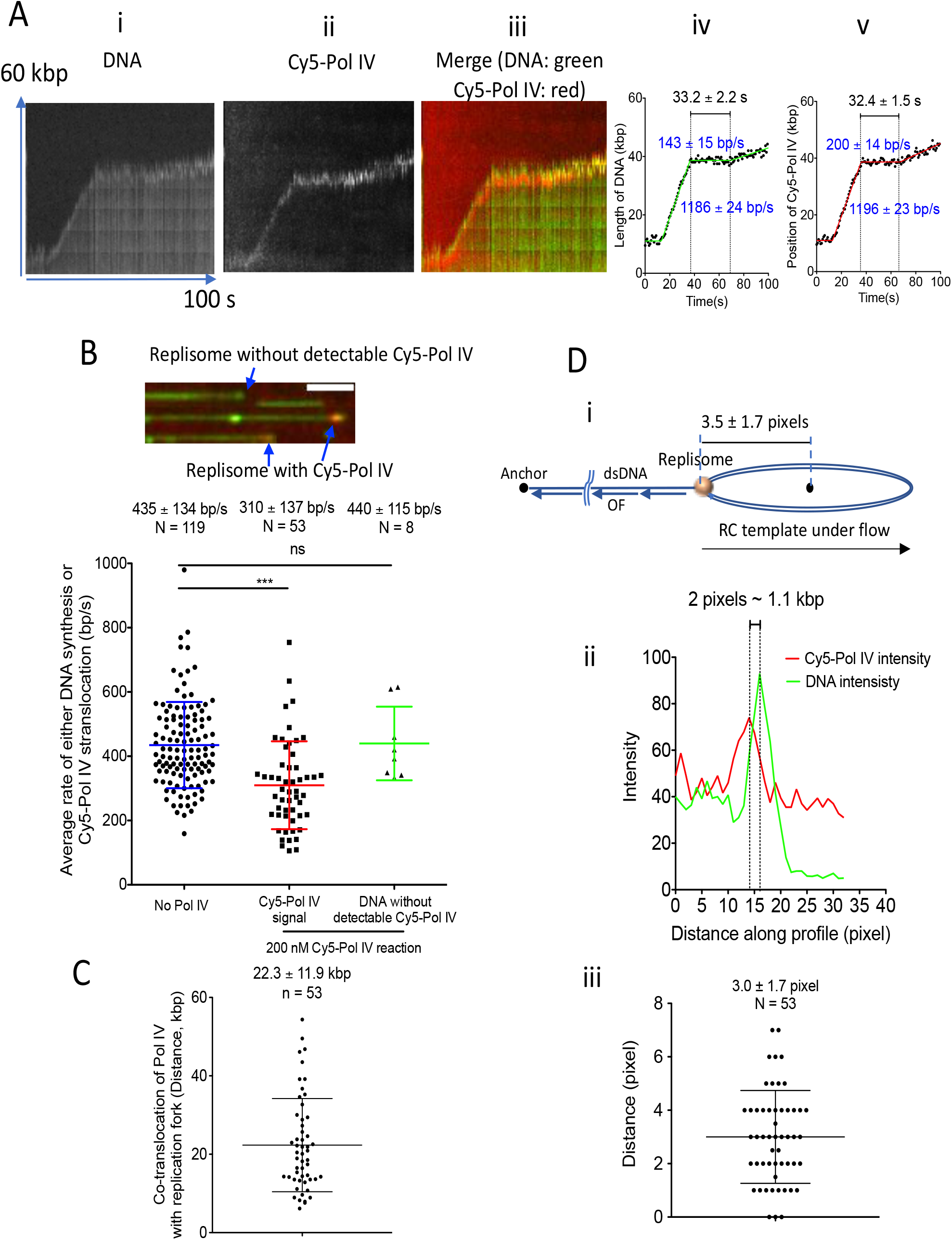
Pol IV localizes to the replisome during replication fork progression. (A) Representative kymographs and tracking trajectories from a reaction where 200 nM Cy5-Pol IV was included in the assembly reaction. (i), (ii), (iii), (iv), (v), SYTOX kymograph (DNA), Cy5-Pol IV kymograph, merge of (i) and (ii), tracking trajectory of DNA length, and tracking trajectory of Cy5-Pol IV signal, respectively. (B) Top panel, expanded merged microscope frame of SYTOX and Cy5 signals showing actively replicating templates either with and without Cy5-Pol IV signals (10%) at the template end. Scale bar is 5 μm. Bottom panel. Average rate of Cy5-Pol IV translocation and the average rate of replication fork progression of DNA molecules without Cy5-Pol IV. The average rate of replication forks formed in the absence of Pol IV is reproduced from Figure 1E. (C) Observed extent of Cy5-Pol IV co-translocation with the SYTOX replicating DNA signal. (D) Colocalization of Cy5-Pol IV and SYTOX template signals. (i) A cartoon showing the expected distance from center of the template signal to the position of the replisome when the system is under flow. (ii) Representative profiles of the Cy5-Pol IV (red) and DNA signals (green) at identical positions in both channels. (iii) Average distance from the center of the Cy5-Pol IV signal to the center of SYTOX template signal. Mean values are given with standard deviations. N, molecules.

These experiments are conducted under flow. As can be seen in the supplemental movies, the circular template, which appears as a bright spot, moves away from the anchored 5’single-stranded end as replication proceeds and is always at the end of the extended DNA. This places the replisome at the junction between the circular template image and the extended double-stranded DNA (Figure 3D). The circular DNA template itself is also extended by the flow into an elliptical shape which appears as a diffraction-limited line of ~7 pixels when fully extended (Figure 3D). Thus, one would expect the replisome to be present on the trailing edge of the extended circular template signal (Figure 3D-i). In our experiments, the center of the Cy5 signal was either co-incident with or always behind the center of the DNA template signal (the tracking algorithm tracks the centroid of the DNA template signal). As an example, in the kymograph shown in Figure 3A, the distance between the two signals was 2 pixels (Figure 3D-ii). The long dimension of the elliptical template under flow was 3.5 ± 1.7 pixels (Figures 3D-i and S3A) and the average distance from the center of the Cy5 signal to the center of the circular template signal was 3.0 ± 1.7 pixels (Figure 3D-iii), thereby locating Pol IV within 0.4 pixels (roughly 200 bp) of the position of the replisome, a value lower than the limit of resolution in our experiments. We therefore conclude that Pol IV assembles with the replisome and moves along with it for long distances that reflect the intrinsic processivity of replication fork progression.

### Co-residence of Pol IV and Pol III on β likely accounts for slowing of the replisome

Pol IV binding to β occurs through two mechanisms: the C-terminus of Pol IV contains a canonical β clamp binding domain (37), an alpha helix that interacts with interdomain connecting loops on the clamp, and also a region of Pol IV known as the “little finger” domain binds to the rim of the β torus (38). Deletion of five C-terminal amino acids of Pol IV inactivates binding to β. We determined whether this mode of Pol IV β binding was required for the observed slowing of replisome progression. This proved to be the case.

The presence of 200 nM Pol IVDC5 in the “Assembly” step of the rolling circle reactions had no effect on either the average replication fork rate (Figure 4A), processivity (Figure 4B), or DNA synthesis burst rate (Figure 4C) of replisomes indicating that, at a minimum, initial association of Pol IV with the replisome is via interaction with β.

**Figure 4:**
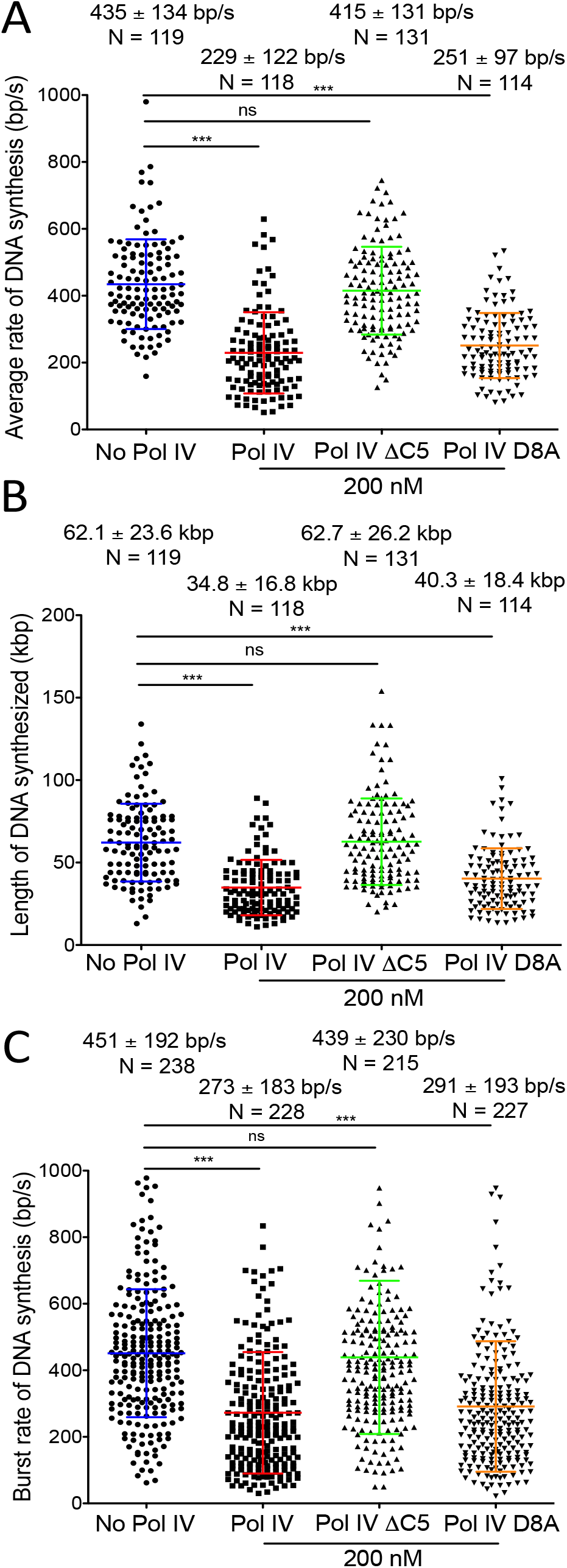
A catalytically inactive Pol IV slows replisome progression. (A) Average rate of replication fork progression from live imaging reactions in the presence and absence of mutant Pol IV proteins as indicated in the assembly reaction. Data for reactions assembled in the absence of Pol IV and in the presence of wild-type Pol IV are reproduced from Figure 1E. (B) Processivities of replication forks from the live imaging reactions in the presence of the indicated concentrations of Pol IV mutants in the assembly step. Data for reactions assembled in the absence of Pol IV and in the presence of wild-type Pol IV are reproduced from Figure 1G. (C) Average burst rates of DNA synthesis in the presence or absence of the indicated concentrations of Pol IV mutants in the assembly step. Data for reactions assembled in the absence of Pol IV and in the presence of wild-type Pol IV are reproduced from Figure 2C. Mean values are given with standard deviations. (*) and (***) denote p < 0.05 and p < 0.0001, respectively, determined by the student’s t-test. N, molecules.

To assess whether Pol IV catalytic activity was required for Pol IV slowing of the replisome, we used the Pol IV D8A variant that is incapable of catalyzing DNA synthesis (39). Surprisingly, 200 nM Pol IV D8A was as active as wild-type Pol IV in slowing the average rate of replication fork progression (Figure 4A), processivity (Figure 4B), and DNA synthesis (Figure 4C).

We demonstrated that Pol IV forms a long-lasting complex with the replisome in a fashion dependent on β clamp-binding that results in a decrease in both the average rate of replication fork progression and the average rate of DNA synthesis bursts, consequently, we conclude that the equivalent effects of Pol IV D8A suggest that Pol IV achieves replisome slowing by being co-resident with the leading-strand Pol III on the same β dimer. An alternative possibility, which we consider unlikely, is that Pol IV, by binding to additional subunits of the Pol III HE, influences the chemical rate of Pol III DNA polymerization directly.

### Pol IV association with the replisome is stabilized by the presence of the *γ* subunit of the Pol III HE

Whereas whether the Pol III HE has two or three α subunits in vivo is problematic, it is the case that E. coli cells engineered to eliminate the chromosomal translational frame shift in *dnaX* that produces the γ subunit of the DnaX complex (40, 41) and expressed a C-terminally tagged γ subunit from a plasmid, yield Pol III HE with one copy of tagged γ; whereas cells that express only τ become sensitive to UV-irradiation and have reduced mutagenic break repair (7). Because of this genetic connection, we asked whether there was any difference in the effect of Pol IV on replisomes formed with Pol III HEs containing two (DiPol) or three (TriPol) copies of the α subunit.

Interestingly, whereas TriPol replisomes were slowed by the presence of Pol IV (Figure 5A), higher concentrations of Pol IV were required to achieve equivalent reductions that were observed with DiPol replisomes (Figure 5B). This was most apparent at 500 nM Pol IV, where TriPol replisomes were still capable of replication (Figures 5A and 5B), whereas no replication was observed with DiPol replisomes (Figures 5B and Table 1). As with DiPol replisomes, Pol IV was slowing the average rate of fork progression of TriPol replisomes by reducing the average rate of DNA synthesis (Figure 5C). These data suggest that the presence of g in the replisome stabilizes its association with Pol IV.

**Figure 5:**
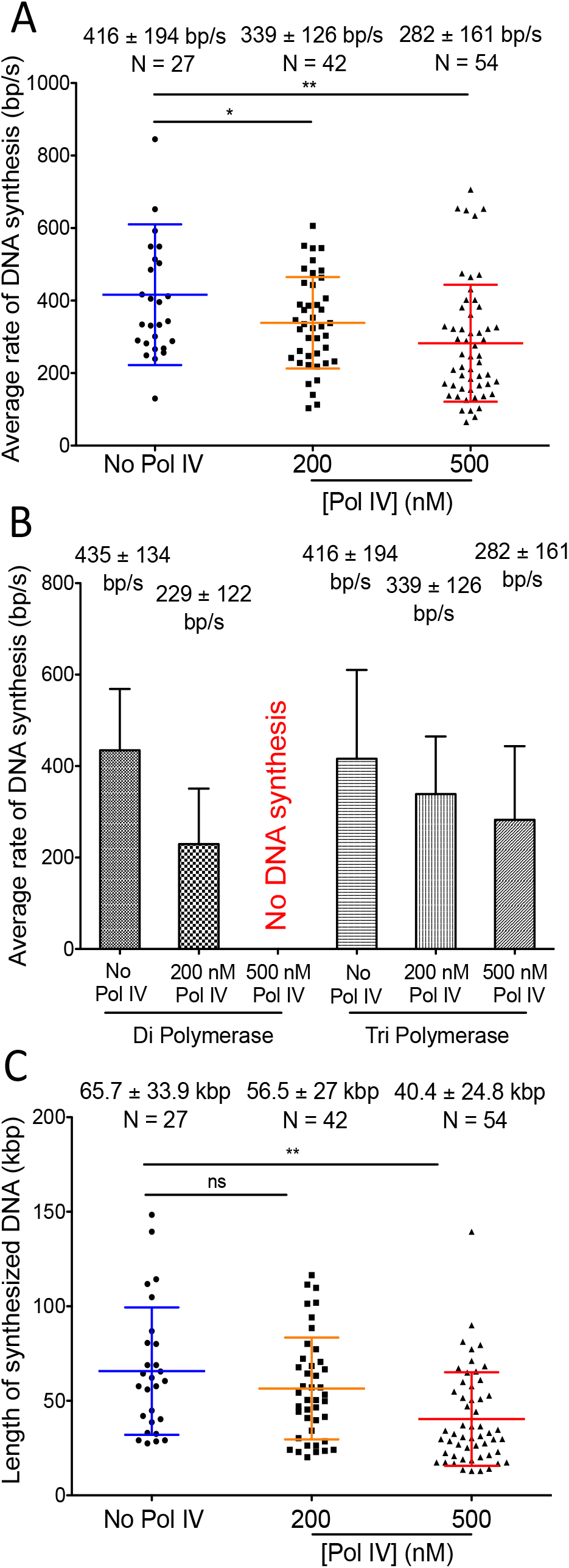
Pol IV is less effective in slowing TriPol III HE replisomes than it is with DiPol III HE replisomes. (A) Average rates of replication fork progression from live imaging reactions with the Tri-Pol III replisome in either the absence of Pol IV or in the presence of the indicated concentrations of Pol IV in the assembly step. (B) Comparison of average rate of replication fork progression between DiPol III HE replisomes and TriPol III HE replisomes in either the absence of Pol IV or in the presence of the indicated concentrations of Pol IV in the assembly step. Data for the DiPol III HE replisomes is reproduced from Figure 1E. (C) Processivities of TriPol III HE replication forks from the live imaging reactions in the presence or absence of the indicated concentrations of Pol IV in the assembly step. Mean values are given with standard deviations. (*) and (***) denote p < 0.05 and p < 0.0001, respectively, determined by the student’s t-test. N. molecules.

## Discussion

Our salient findings are that TLS DNA polymerase IV forms a stable complex with Pol III HE replisomes during their assembly onto primer-template DNA and that this association persists during DNA replication for the apparent lifetime of the replisome, causing the average speed of the replisome to decrease by about half. These findings have interesting ramifications for replication fork progression under conditions of cellular stress, the manner by which Pol IV and Pol III associate during DNA replication with the β clamp, the mechanism of polymerase switching, and the manner by which Pol IV surveilles DNA template damage.

### Association of Pol IV with the replisome and polymerase switching

The concept of a TLS and replicative polymerase binding to the same β dimer in an active replisome is an attractive one, providing the replisome with a “tool belt” to fix template problems by switching between polymerases as it moves along, without undue slowing of the overall replication process (29). This model was premised to a large extent on the fact that the E. coli TLS DNA polymerases possessed canonical β clamp binding motifs as did the a subunit of the Pol III HE thus, in principle, allowing one β clamp dimer bound to the DNA to bind two different DNA polymerases. The idea that two DNA polymerase could simultaneously occupy one β dimer became somewhat complicated by the observation that in a cryo-electron microscopy structure of TriPol III HE bound to a primer-template, both binding sites of one β were occupied: one by the β-binding motif on an α subunit, and one by a weaker β-binding motif on the ε subunit (42).

Polymerase switching has been investigated by several groups using several different approaches. The actual switching event is often scored by taking advantage of the dramatic difference in the rate of polymerization between Pol IV and the Pol III HE on primed singlestranded templates in either ensemble or various single-molecule types of experiments. For example, Heltzel et al. (30) showed that the little finger domain of Pol IV was not required for Pol IV polymerase activity but was required for polymerase switching and that only one binding site on β was sufficient to observe the switch. Subsequent single-molecule experiments supported these observations and led to a model where initial contact between Pol IV and β bound to Pol III occurred in a concentration-dependent manner via the little finger domain leading to subsequent competition between Pol IV and the ε subunit of the Pol III HE for one of the β binding sites (43). Similar experiments demonstrated switching between Pol II and Pol III as well as isolation by gel filtration of a ternary complex of β, Pol II, and the Pol III core (44).

On the other hand, using co-localization single-molecule spectroscopy of fluorescently-labeled Pol IV and Pol III, Zhao et al.(45) have argued that neither Pol II nor Pol IV form a stable complex with Pol III on β loaded to a primer-template, but alternate binding to the clamp. Similarly, Furukohri (39) et al. demonstrated in ensemble experiments that Pol IV could displace a stalled Pol III HE from a primer-template, and argued that switching was the result of the complete exchange of Pol IV and Pol III on β. Interestingly, these authors also found that the Pol IV variant Pol IV D8A DC5, which is catalytically inactive and does not bind to β via the clamp-binding motif, also promoted Pol III dissociation from β, suggesting the presence of another interaction between Pol IV and the Pol III HE. As we have demonstrated, TriPol Pol III HE, which lacks the γ subunit, is less sensitive to replisome slowing by Pol IV than DiPol Pol III HE. It is therefore possible that this second interaction between Pol IV and the Pol III HE may be with the DnaX complex.

In our studies, we assembled replisomes in the presence or absence of Pol IV, removed any Pol III HE and Pol IV that were not bound to the template, and assessed subsequent replisome-catalyzed replication fork progression. The observed reduction of replisome speed by Pol IV was concentration-dependent and dependent on the presence of the β clamp-binding motif. However, this does not necessarily mean that the initial association of Pol IV with the replisome during assembly was via β clamp binding, but if it was not, it does imply that whatever other interaction operated, it had to be fairly stable to resist washing. Within the limits of experimental error, we show that Pol IV is associated with the replisome and remains associated, as measured by the co-incident processivities of replisome signal (the SYTOX Orange-stained DNA trajectory) and fluorescent Pol IV trajectory, for the lifetime of the replisome, further indicating that the association between Pol IV and the replisome was quite stable.

Only replisomes that had bound Pol IV exhibited slowing of replication fork progression. Clearly, this observation indicates that the bound Pol IV interfered in some manner with polymerization by Pol III. Furthermore, the manner of this interference was not dependent on additional molecules of either Pol IV or Pol III in solution, the phenomenon of replisome slowing was manifested by the action of one Pol III HE particle and one Pol IV molecule within a moving replisome. Importantly, Pol IV catalytic activity was not required for replisome slowing. These observations present only a few possible mechanisms to account for replisome slowing: 1) Pol IV loosely bound to the rim of β, or possibly to the DnaX complex, exerts an allosteric effect on the active site of a that alters the speed of polymerization. We know of no data that would support this possibility. 2) Pol IV bound to the DnaX complex interacts with the DnaB helicase, which is also bound to the τ subunit of the DnaX complex (46), and slows template unwinding. This possibility is quite unlikely because it is the case that the polymerase pushes the helicase (23, 47) 3) Pol IV residence on the same β clamp to which the leading strand α subunit of the Pol III HE is bound, with each polymerase occupying one binding cleft on β, directly results in replisome slowing (Figure 6B). And 4) Pol IV and a are rapidly and continuously switching on the β clamp, resulting in an overall decrease in the rate of DNA synthesis (Figure 6C). We believe that possibility three is the most likely explanation for our results.

**Figure 6:**
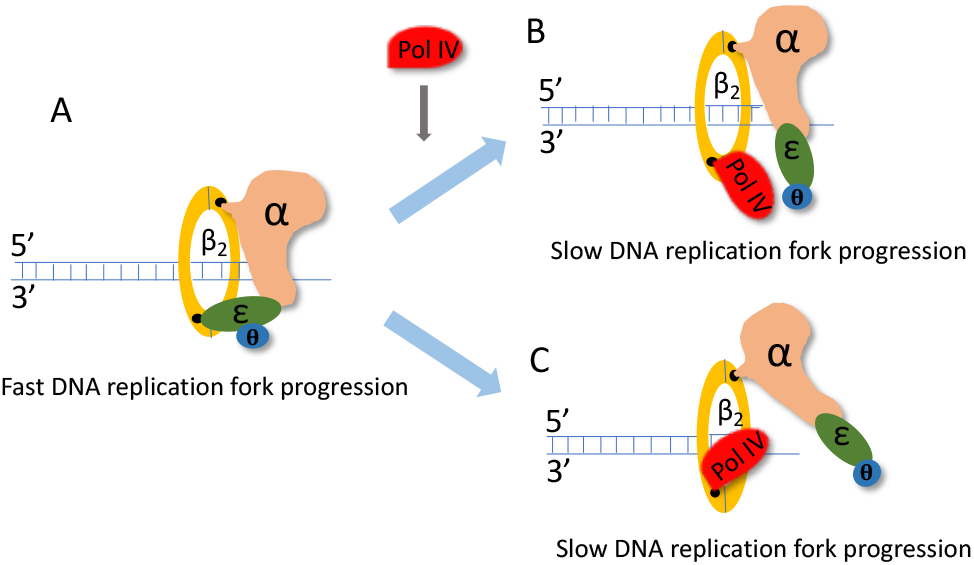
Association of Pol IV with the leading-strand β clamp slows the rate of replication fork progression. (A) Fast replication fork progression in the absence of Pol IV. (B) Pol IV completes with the ε subunit of the Pol III HE for one of the binding clefts on the β dimer. (C) Pol IV associated with the leading-strand β clamp competes directly with the leading-strand α subunit of the Pol III HE for the active 3’-OH of the nascent DNA.

In our previous studies on replication fork progression (23), we showed that β bound to the leading-strand polymerase did not have an infinite residence time in complex with the replisome. In reactions where only leading-strand synthesis was observed (primase was omitted from the experiments), the absence of β in the flow resulted in a decrease in average processivity of about 50% and, tellingly for our observations reported herein, generated a double Gaussian distribution of DNA synthesis burst rates, one slow, at 270 nt/s, and one fast, at 560 nt/s. We speculated at the time that perhaps some molecules of Pol III HE had lost some contact with β, thereby generating the slower moving population of replisomes. We suggest this phenomenon provides the most reasonable explanation for the effect of Pol IV on replisome progression. We believe it likely that Pol IV competes with the ε subunit of the Pol III HE for the second binding cleft on β, and that the lack of two contacts between the Pol III HE and the β clamp results in the overall decline in the rate of DNA synthesis. A catalytically inactive Pol IV would still retain replisome-slowing activity in this scenario because it would be expected to bind β. We cannot rule out possibility four, rapid switching between the two polymerases on the active 3’-end of the primer, but it is not necessary to explain our data, whereas possibility three is sufficient. Furthermore, the time scale of such rapid switching, if it occurs, is far beyond the time resolution of our experiments.

### Template surveillance during the SOS response

The replisome slowing activity of Pol IV that we have observed is concentration dependent. A nearly full extent of the effect was observed at 200-300 nM Pol IV, consistent with the observations that overproduction of Pol IV in vivo under stressed (27) and unstressed (25, 26) conditions, even in the absence of template damage, slows fork progression and with initial estimates of the concentration of Pol IV in the cell in the presence and absence of the SOS response at 250 copies per cell (19). More recent data, however, pegs Pol IV concentration in cells to be less than one-tenth of previous estimates: 6 nM in unstressed cells and 34 nM in SOS-induced cells (20), suggesting that Pol IV should not associate with the replisome under any condition. Yet the same authors do observe fluorescent Pol IV co-localizing with fluorescently labeled τ and ε subunits of the Pol III HE. Whereas there is no obvious answer to this conundrum, it may be that molecular crowding in the nucleoid of E. coli cells increases the effective concentration of Pol IV, allowing it to associate with the replisome when SOS is induced.

A replisome that carries both Pol IV and Pol III continuously during DNA synthesis doesn’t seem like a good idea under normal circumstances, given the obvious increased potential for polymerase switching. Afterall, TLS DNA polymerases are error prone (11) and such a scenario should increase the overall rate of mutagenesis. However, it is possible that Pol IV does not switch with Pol III unless a template lesion is encountered. Thus, it might be the case that Pol IV does double duty to help overcome large amounts of template damage encountered during the SOS response: slowing replication forks to decrease the probability of encounters with template lesions and surveilling the template as the replisome progresses, bypassing any template damage that is encountered.

## Materials and methods

### Proteins

Replication proteins were as described previously (23). Pol IV protein was purified from 4 L BL21(DE3) pLysS (pET16b-*dinB*) (48) that had been grown in LB medium to O.D._600_ = 0.7 at 30 °C and induced for 4 h at 30 °C by the addition of I mM IPTG. Cell pellets were resuspended in Lysis buffer (50 mM Tris-HCl (pH 7.5), 1 M NaCl, 10% sucrose, 2 mM DTT, 1 mM EDTA, and 1X protease Inhibitor cocktail (Roche)), lysozyme was then added to final concentration of 2 mg/ml, the suspension stirred for 30 min at 4 °C, followed by 5 min at 37 °C for 5 min, and the suspension centrifuged at 15,000 x g for 30 min. Protein was precipitated from the cleared lysate by the addition of (NH_4_)SO_4_ to 30% saturation, collected by centrifugation, resuspended in Heparin column binding buffer (50 mM Tris-HCl (pH 7.5), 10% glycerol, 10 mM NaCl, 2 mM DTT, 1 mM EDTA), and loaded onto a HiTrap Heparin column (Pharmacia) equilibrated with the same buffer. Pol IV was eluted with a gradient of 10 mM to 1 M NaCl in the same buffer.

Pooled Pol IV fractions (identified by SDS-PAGE) were dialyzed against MonoQ column binding buffer (50 mM Tris-HCl (pH 7.5), 10% glycerol, 100 mM NaCl, 2 mM DTT, 1 mM EDTA) overnight and loaded onto a MonoQ column equilibrated with the same buffer. Pol IV, which passed through the column, was dialyzed against storage buffer (50 mM Tris-HCl 7.5, 10 mM NaCl, 1 mM EDTA, 2 mM DTT, 10% glycerol) and stored at −80 °C.

His-tagged Pol IV proteins (HT-Pol IV, HT-Pol IV ΔC5 and HT-Pol IV D8A mutants) were purified as described previously (24) with the exception that a Heparin column as described above was used as the last step instead of gel filtration.

### Pol IV DNA polymerase assay

The DNA template used to measure the DNA polymerase activity of Pol IV and mutants was prepared by annealing a ^32^P-labeled 25-mer oligonucleotide labeled with ^32^P at the 5’-end, to circular M13 ssDNA. Reaction mixtures (20 μl) containing 20 mM Tris-HCl (pH 7.5), 8 mM MgCl2, 5 mM DTT, 0.1 mM EDTA, 40 μg/ml BSA, 4% glycerol, 200 μM each dNTPs, 1 μM SSB, 8 nM primed template (5’-^32^P-CGACGTTGTAAAACGACGGCCAGTG-3’annealed to M13 Ophrys ssDNA (8.6 knt) (49)) and the indicated concentrations of Pol IV were incubated at 37 °C for 5 min. Reactions were stopped by the addition of a 2-fold volume of loading buffer (50 mM EDTA, 1 mg/ml xylene xyanol FF, and 1 mg/ml Bromophenol Blue in formamide). Samples were denatured at 95°C for 5 min and the DNA products analyzed by denaturing gel electrophoresis (12% polyacrylamide (acrylamide:bisacrylamide 29:1), 8 M Urea, and 1 X Tris-borate-EDTA buffer (TBE, 89 mM Tris base, 2 mM EDTA, 89 mM boric acid) at 8 V/cm for 2 h at room temperature. The gel was dried onto Whatman paper filter (GE Healthcare Life Sciences) at 80 °C under vacuum for 2 hours. Radiolabelled products were scanned using a STORM 860 PhosphorImager (Molecular Dynamics). The intensity of unextended primer (^32^P-labeled 25-mer oligonucleotide) was quantified by ImageQuant 5.2 software.

### Cy5-labeling of Pol IV

Cy5-Pol IV was prepared using Cy5-NHS ester (Sigma) as previously described (50). Pol IV (100 μg) was added to 100 μl of buffer L (50 mM K_2_HPO_4_/KH_2_PO_4_ (pH 7.0), 100 mM NaCl, 0.1 mM DTT, and 10% glycerol) followed by the addition of a 5-fold excess of Cy5-NHS (20 mM in DMSO). The mixture was incubated at 4 °C for 4 h in the dark. Free fluorescent dye was removed by filtration through a P10 (Bio-Rad) column equilibrated with the same buffer. Fractions were collected (60 μl) and Pol IV and free Cy5 were measured by following O.D._280_ and O.D._650_ using a Nanodrop spectrophotometer. The degree of Pol IV labeling was determined by measuring O.D._280_ and O.D._650_. The extinction coefficient of Pol IV is 79,700 M^-1^ cm^-1^ and the extinction coefficient of Cy5 is 250,000 M^-1^ cm^-1^. Concentrations of Pol IV and Cy5 were calculated as below:

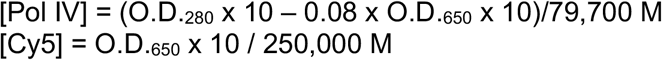

### Single-molecule rolling-circle replication reaction

The preparation of the rolling circle template, cover glass, and flow-cell were done as described previously (23). The inlet line to the flow cell was designed with two loops, one for the “Assembly” step and the other for the “Start” step (Fig. S1A). The “Assembly” loop was filled with 50 μl of “Assembly” reaction (75 nM SYTOX Orange, 60 nM DnaB_6_, 380 nM DnaC810, 20 nM Pol III* (or the same concentration of TriPol*, as indicated), 30 nM β_2_, 40 μM each of 3 dNTPs, 200 μM NTPs, 100 μg/ml BSA, and Pol IV or Cy5-Pol IV as indicated in SMB (50 mM HEPES-KOH, pH 8.0, 15% sucrose, 25 mM KCl, 10 mM Mg(OAc)_2_, and 50 mM DTT). The “Start” loop was filled with 150 μl of “Start” reaction (75 nM SYTOX Orange, 320 nM DnaG, 30 nM β_2_, 1 μM SSB, 40 μM dNTPs, 200 μM NTPs, and 100 μg/ml BSA in SMB buffer). The reaction was started after pushing 100 μl of solution through the lines at a flow rate of 1250 μl/h. Elongation reactions were recorded for 7 min at a frame rate of 7 Hz using a DU-897E iXon EMCD camera (Andor).

The flow cell was mounted on an Eclipse TE2000-U inverted microscope (Nikon) with a TIRF attachment and imaged using a CFI Plan Apo TIRF 100X, 1.45 numerical aperture, oil-immersion objective as described (23). Double-stranded DNA stained with SYTOX Orange (Ex/Em 547/570) and Cy5-Pol IV (Ex/Em 649/670) were visualized by continuously illuminating the sample with 561 nm and 640 nm lasers at about 70 μW. Light was collected with a Dual View cassette (Optical Insights) where the green and red signals were separated by a DAPI/Cy5 dichroic mirror, followed by emission filters ET605/70 (Chroma) and ET524/40-665/80 (Chroma) to simultaneously visualize double-stranded DNA and Cy5-Pol IV signals.

The length of synthesized DNA was measured manually and analyzed using ImageJ (NIH, v. 1.52). Registration of dual view images was calibrated every day by using 1 μm beads labeled with Cy5 and Cy3.

### Analysis of the replication fork progression in the live reactions

To analyze the progression of replication forks in the live reaction, raw images were averaged using a custom-written ImageJ plug-in (23), a 5-frame sliding window, that reduced the stack size by a factor of five and defined the time resolution to about 700 ms. Molecules of interest were cropped from the averaged images and analyzed individually. Gaussian Blur, an averaging filter in ImageJ v.1.52k that uses convolution with a Gaussian function, was applied for smoothing and minimizing the background noise of the images. Subsequently, the images were binarized using the Auto-Threshold function in ImageJ. DNA length in each frame of a time course was measured using a custom-written ImageJ plug-in (Ichiro Amitani and Katsumi Morimatsu) that detects the edges of the extended molecule. The tracked length of DNA in pixels was converted to base pairs using a calibration described previously (23). Plotting the tracking DNA length (kbp) and time (s) in Prism 5.0b yielded the trajectory of replication fork progression. Trajectories were fitted by a nonlinear regression method using multiple segment lines constraining either the end (3-segment fitting) or both the end and middle segments (5-segment fitting) to zero (23). Rates of replication fork progression were determined by dividing the length of extended DNA by the time elapsed between the initiation and termination of DNA synthesis. DNA length at the end point of a trajectory was assigned as the processivity of replication fork progression. Burst rates of DNA synthesis and pause times were derived from the line segment fitting.

### Analysis of the distribution of replication fork rates

The distribution of DNA synthesis rates in absence of Pol IV were analyzed as the sum of two Gaussian distributions using Prism (V.9.3.1). To enable convergence, the mean value and standard deviation (SD) of the distribution in the absence of Pol IV were used to constrain the parameters of one Gaussian distribution (406 ±117 bp/s). Fitting of the 200 nM Pol IV distribution defined the second, slower distribution as 185 ±90 bp/s. These parameters were used to simulate double-Gaussian distributions for the remaining rate profiles in the presence of Pol IV. The relative amount of each component was calculated from the relative amplitude of each Gaussian distribution.

### Analysis of the Cy5-Pol IV translocation in live reactions

Images of Cy5-Pol IV live reactions were averaged as above. Merging the kymographs for SYTOX Orange-stained dsDNA and Cy5-Pol IV that were at the same coordinates in the imaging channels created the composite image kymograph. The individual Cy5-Pol IV signals were tracked by using a custom-written ImageJ plug-in (Spot Track3, Ichiro Amitani). The tracking trajectories were fitted with multiple-segment lines to analyze the rate and distance of Cy5-Pol IV translocation.

Co-localization of Pol IV and the replisome was determined based on the distance from the Cy5-Pol IV signal to the center of the template signal at the end of the replicating DNA. A onedimensional line was drawn manually across a DNA molecule in the SYTOX Orange-stained DNA channel and the Plot Profile function in ImageJ was used to generate its intensity profile. The center of the template signal was assigned as the peak of fluorescence intensity along the profile line. The same line coordinates were used to generate the Cy5-Pol IV intensity profile and the position of Cy5-Pol IV was assigned as the peak of the Cy5 intensity profile. The distance between Cy5-Pol IV and the center of the template was determined as the distance between these two peaks of the intensity profiles.

## Supporting information

Supplemental Movie S2B(iii)

## Acknowledgments

We thank Soon Bahng (MSKCC) for purified proteins, Dr. Asako Furukohri (NAIST) for the Pol IV mutant plasmids, and Dr. Phuong Pham (USC) for the native Pol IV strain. We also thank all members of the S.C.K. group, especially James Graham and Neville Gilhooly for advice and Ichiro Amitani and Katsumi Morimatsu for custom written analyses programs. These studies were supported by NIH grants R35 GM126907 to K.J.M., NCI Cancer Center Support Grant NCI P30CA008748 to MSKCC, and R35 GM131900 to S.C.K.

## Supplementary Information

### Supplementary Figure and Movie Legends

**Figure S1:**
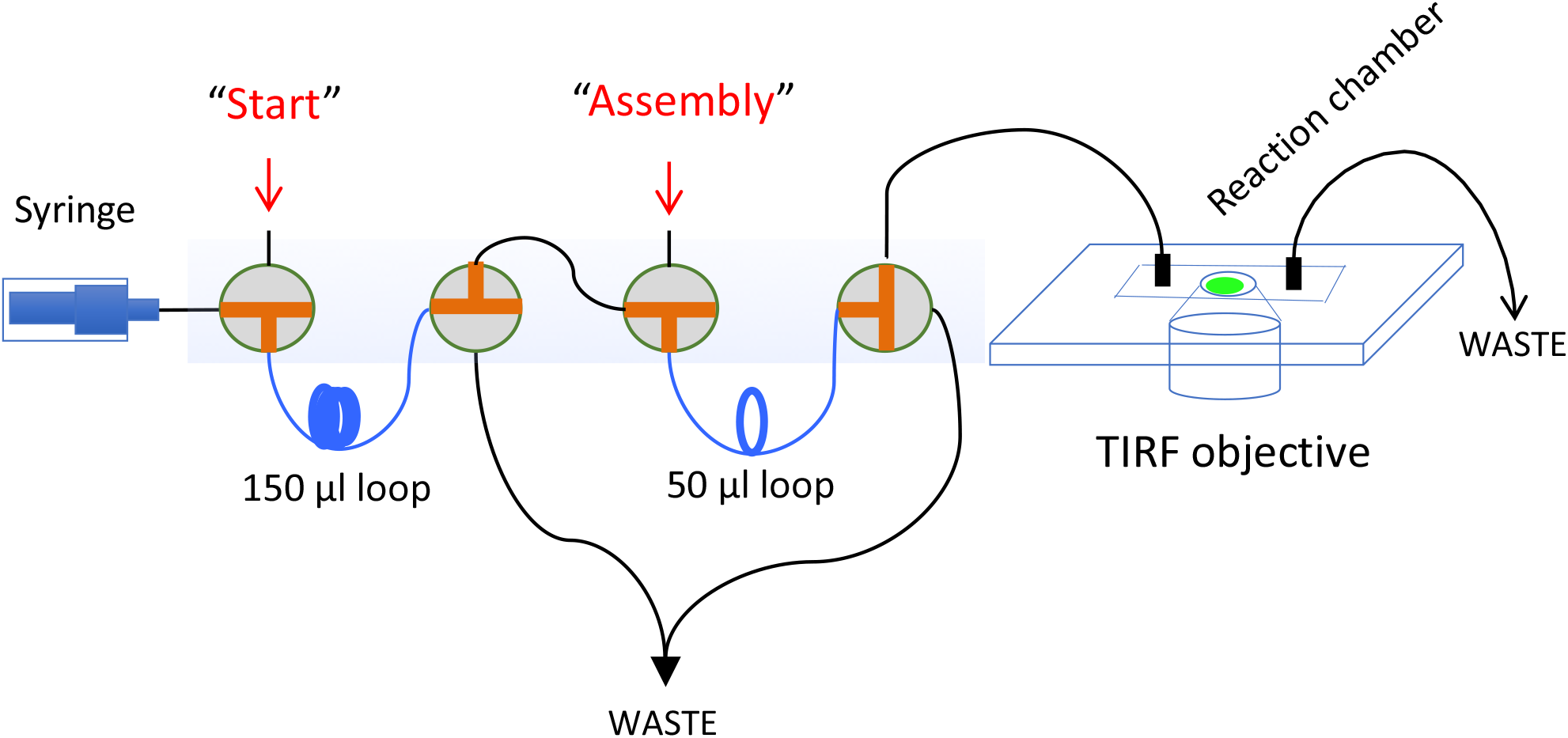
Schematic representation of TIRF single molecule assay, related to Figure 1. The Assembly and Start reactions are loaded into 50 μl and 150 μl loops, respectively. The pump pushes the reactions to the chamber at a flow rate 1250 μl/hr.

**Figure S2:**
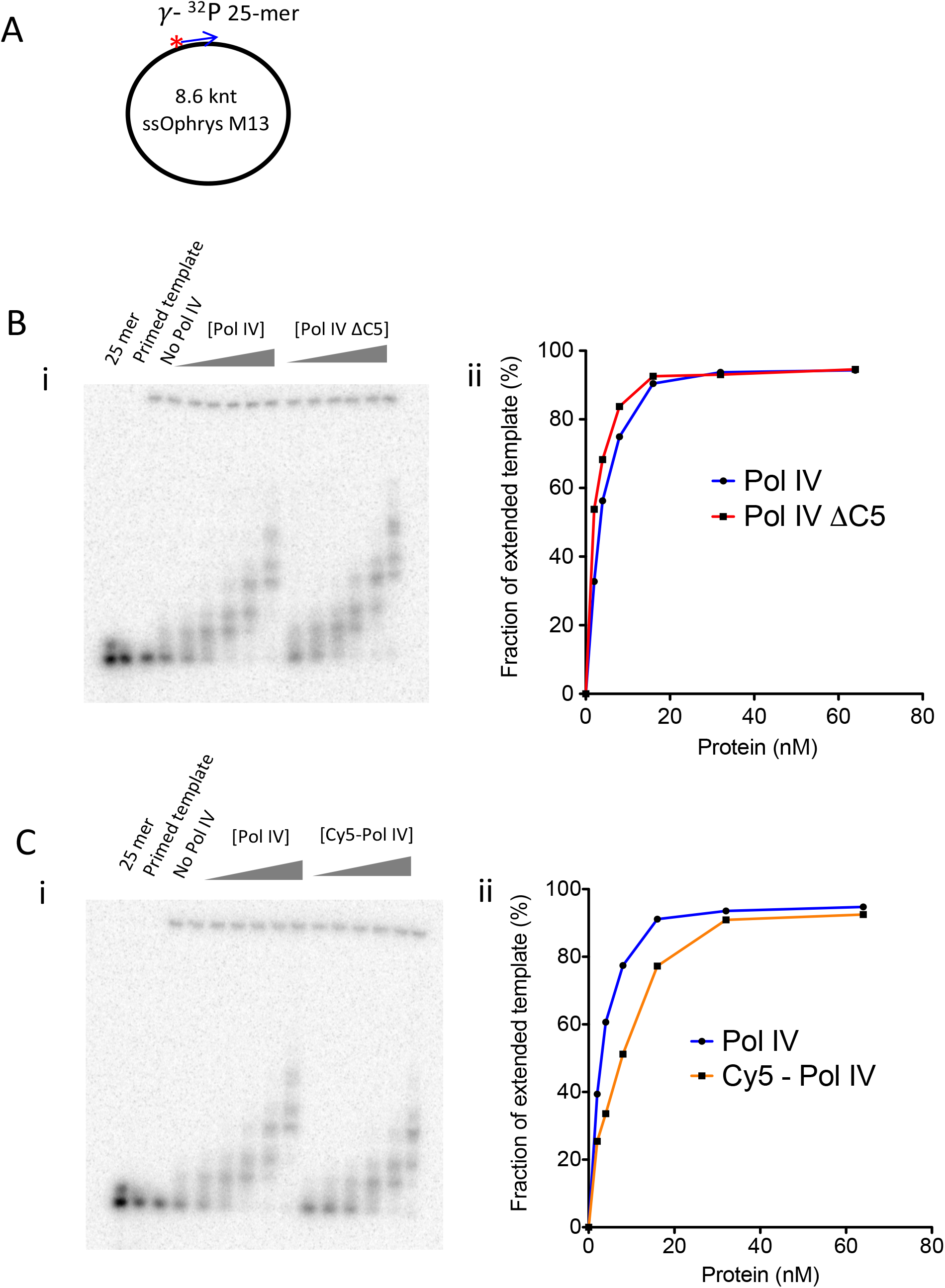
Activity of Pol IV, Pol IV ΔC5 and Cy5-Pol IV, related to Figure 3 (A) Primer/Ophrys M13 DNA template. A ^32^P-labeled 25-mer primer annealed to 8.6 knt Ophrys M13 circular ssDNA. (B) Extended products in presence of Pol IV and Pol IV ΔC5 (i) at 0, 2, 4, 8, 16, 32, 64 nM. (ii) Graph showing fraction of extended products. (C) Extended products in presence of Pol IV and Cy5-Pol IV (i) at 0, 2, 4, 8, 16, 32, 64 nM. (ii) Graph showing fraction of extended products.

**Figure S3:**
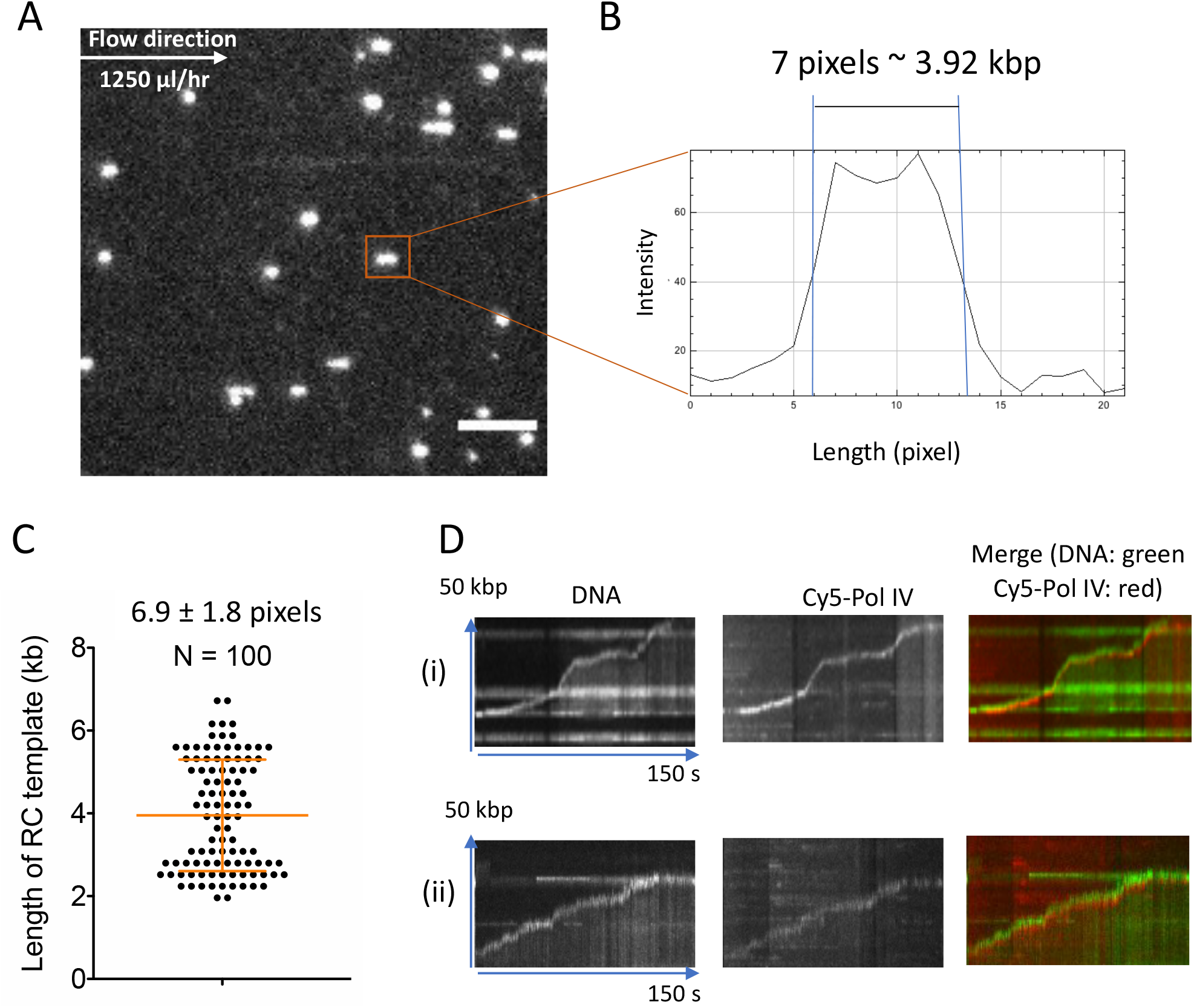
Determining the distance from center of RC DNA template to the replisome, related to Figure 3. (A) Typical image of RC DNA template stained by SYTOX Orange on the glass surface under flow before loading the Assembly and Start reactions. Scale bar is 5 μm. (B) The length of RC DNA template is determined by using intensity profile of the molecules. (C) Average length of RC template under the flow. (D) (i) (ii) Kymographs of two typical molecules from two independent reactions showing live imaging of Cy5-Pol IV.

Movie S1A. Direct visualization of replication in the absence of Pol IV, related to Figure 1. Movie of the same field as Figure 1B. The red “x” marks the molecule analyzed in Figure 1C. Scale bar, 5 μm, equivalent to 17.5 kb dsDNA.

Movie S1B. Cropped movie of the molecule is marked as the red “x” in Movie S1A. Scale bar, 5 μm, equivalent to 17.5 kb dsDNA.

Movie S2A. Direct visualization of replication in the presence of 200 nM Pol IV in the Assembly reaction, related to Figure 2. The red x marks the molecule analyzed in Figure 2A and B, iv. Scale bar, 5 μm, equivalent to 17.5 kb dsDNA.

Movies S2B i-v. Movies of the molecules shown in Figure 2A i-v, respectively. Scale bar, 5 μm, equivalent to 17.5 kb dsDNA.

Movie S3. Direct visualization of Cy5-Pol IV at the replisome, related to Figure 3. Composite image is made by overlaying both color channels (DNA is green and Cy5-Pol IV is red). Scale bar, 5 μm, equivalent to 17.5 kb dsDNA.

## Notes

### Competing Interest Statement

The authors have declared no competing interest.

